# Ascorbic acid and gallic acid mitigate behavioural deficits and loss of cerebellar myelin induced by Bisphenol-A in Wistar rats

**DOI:** 10.1101/2024.07.17.603899

**Authors:** Kabirat Oluseun Adigun, Michael Ibukun Ehinmowo, Oluwabusayo Folarin, Olamide Elizabeth Adebiyi

## Abstract

**Background:** Bisphenol-A (BPA) is a chemical component of plastics, polycarbonates, and epoxy resins making it an environmentally ubiquitous chemical. Its toxicity has been linked to several disorders including neurological disorders. Gallic acid (GA) and Ascorbic acid (AA) are both antioxidants that have been reported to have neuroprotective effects. We therefore investigated the neuroprotective effects of GA and AA on BPA-induced neurotoxicity.

Forty-eight (48) Wistar rats were grouped into Control, BPA, BPA+AA, BPA+GA, AA, and GA groups with eight rats in each group. All treatments were administered via oral gavage for 21 days. Behavioral experiments were conducted to assess forelimb strength (hanging wire test), anxiety (open field and elevated plus maze tests), depression (forced swim test), and spatial memory (Morris water maze). Rats were perfused and brain samples were collected for Luxol fast blue staining. Sera were collected to evaluate the levels of inflammatory cytokines via Enzyme-Linked Immunosorbent Assay.

**Results:** There was no significant difference in hanging latency between all the groups. There were also no significant changes in the open field test but the elevated plus maze showed an increase in time spent in closed arms in the BPA group indicative of anxiety. GA and AA reduced the duration of immobility that was elevated in the BPA group which suggests they have antidepressant-like effects. Additionally, we observed disruption in the Purkinje cell layer of the cerebellum in the BPA-alone group.

**Conclusion:** BPA seems to cause anxiety and depressive-like effects which are ameliorated by the gallic acid and ascorbic acid.

## BACKGROUND

Bisphenol-A (BPA 2, 2-bis (40-hydroxyphenyl) propane) belongs to the broad Bisphenols family of chemicals widely used in the synthesis and production of hard and soft plastic products, epoxy resin, and polycarbonate materials [1, 2].

BPA products such as certain cosmetics, toys, dental sealants, plastic food containers, and floorings are predominant in the environment leading to exposure in humans and animals via the oral, respiratory, and transdermal routes [3, 4, 5]. Exposure to BPA can also occur during prenatal development (via intrauterine transmission), early childhood development (e.g., via lactation or direct contact), and adulthood [6, 7]. BPA toxicity has been recognized to be associated with several disorders such as cardiovascular and reproductive dysfunctions, neurodevelopmental disorders, neurodegenerative diseases, infertility, and chronic diseases.

BPA is an endocrine-disrupting chemical that affects endocrine signalling in the body leading to diverse pathological outcomes. Previous studies have shown a direct link between BPA and disease pathogenesis [8, 9]. This is usually via the induction of oxidative stress, that is, by acting on hormone receptors in target cells, BPA’s endocrine signaling function impacts cells and tissues, producing unpleasant physiological responses linked with oxidative stress and inflammation [9]. BPA disrupts redox homeostasis through the increase of oxidative mediators and the reduction of antioxidant enzymes. This results in mitochondrial malfunction, changes in cell signaling pathways, and the induction of apoptosis. As such, BPA in cells and tissues increases oxidative stress due to the elevated production of toxic free radicals [10].

As a result of the effect of BPA on the endocrine system, particularly on its ability to modify the actions of physiologic estrogens, pioneer research primarily investigated its effects on sexual dysfunction, malformation, and cancers of reproductive origin. However, because BPA is lipophilic and can cross the blood-brain barrier, emerging research has shown that BPA also affects neuronal and cortical functions in different life stages and across several animal species. Current research has now established that gestational, perinatal, and neonatal exposures to BPA affect developmental processes, including brain development and gametogenesis, with consequences on brain functions, behavior, and fertility [11].

BPA induces neurotoxicity through several mechanisms including the reduction of synaptic plasticity, inhibition of neurogenesis, generation of oxidative stress, and induction of autophagy and apoptosis [12, 11, 13]. Using animal models, several studies have reported that BPA exposure during the gestational period affects brain development and behaviors. For example, BPA has been linked to alterations in brain structure [14], enhanced depressive-like behavior [15], memory impairment [16], increased anxiety and cognitive deficits [17], disruption of neurotransmitter system [18], impact on social behavior and anxiety [19], and perturbed differentiation and migration of neurons [20].

One of the major contributors to BPA neurotoxicity is the production of free radical species leading to oxidative stress [21, 22], hence we hypothesized that potent antioxidants with neuroprotective effects could ameliorate BPA-induced neurotoxicity. Furthermore, in recent times, there has been a worldwide inclination toward the intake of natural plant-derived antioxidants due to their lower side effects, efficacy in preventing diseases, and general acceptability by the population [23]. Thus, we tested the ameliorative effects of two compounds, known for their anti-inflammatory and antioxidant properties. The first compound, ascorbic acid, is an example of a non-enzymatic antioxidant [24]. Non-enzymatic antioxidants regulate the oxidative environment by interrupting and terminating free radical chain reactions [25]. Ascorbic acid has been reported to have a pro-oxidant effect by which it reduces free radicals to react with oxygen to form a lipid peroxidation generator [26]. Important dietary sources of ascorbic acid for humans and animals include fruits and vegetables. Ascorbic acid protects biomolecules from getting damaged by scavenging oxygen-free radicals in cells [27]. Co-administration of BPA and ascorbic acid for 45 days to male Wistar rats was reported to not affect body and organ weight as the control group given BPA and had a converse effect on hyperchromatic cell number in the brain cortex in increasing the oxidative stress elicited by BPA exposure. Gallic acid (GA) is a trihydroxybenzoic acid with strong antioxidant and free radical scavenging properties [28]. It is one of the most abundant phenolic acids present in various plants and its derivatives can be found in various fruits such as lemon, bananas, pineapples, and some plants [29]. Gallic acid is known to have antioxidant, anti-inflammatory, antimicrobial, anticancer, gastroprotective, cardioprotective, and neuroprotective properties [30]. It has been reported over the years that gallic acid is effective against several neurological and neurodegenerative disorders such as Alzheimer’s, Parkinson’s diseases, depression, and anxiety [31] and traumatic brain injury [32].

## METHODS

### Experimental animals and procedures

Forty-eight adult Wistar rats (male = 24, female = 24) were used for the study. The rats were kept in the Experimental Animal Housing Unit of the University. They were kept in clean polypropylene cages, fed with a standard diet of mice pellets and water, and were allowed to acclimatize to laboratory conditions for 2 weeks before the experiment. This study was approved by the Animal Use and Care Research Committee of our institution.

### Chemicals

The chemicals used in the experiment, including Bisphenol-A (BPA), Ascorbic Acid (AA), and Tween 20 were of analytical grade and were procured from Sigma, USA.

### Experimental design

The animals used in the study were randomly assigned into 6 groups of eight animals each as follows:

- Group A (Control Group): 0.5 ml of Tween 20
- Group B (BPA only Group): 10 mg/kg per day.
- Group C (BPA + Ascorbic Acid Group): BPA (10 mg/kg per day) with a concomitant Ascorbic Acid (100 mg/kg)
- Group D (BPA + Gallic Acid): 10 mg/kg of BPA and 20 mg/kg GA
- Group E (Ascorbic Acid only Group): Ascorbic Acid only (100 mg/kg per day)
- Group F (Gallic Acid only Group): 20 mg/kg body weight GA dissolved in distilled water.

All treatments were administered orally for 21 days.

### Behavioural Tests

#### a. Motor Strength Assessment

##### Forelimb Support (Hanging Wire Test)

A 2 mm thick metallic wire was tightly attached to a frame to avoid vibration or unwanted displacement of the wire during the measurements. Individual rats were placed on the 2 mm diameter hanging wire with their forelimbs and monitored for 2 minutes as described previously [33]. The time it took the animal to stay on the wire before falling was taken and recorded. Rats were rested after the first trial and each rat was allowed to undergo the test twice.

#### b. Tests for Anxiety and Depression

##### The Open Field Maze Test

The open field was used to assess anxiety and locomotion [34]. The open field apparatus was constructed with white plywood and measured 72 × 72 cm with 36 cm walls. One of the walls was covered with plexiglass so the rats could be visible in the apparatus. The area is divided into 25 squares (20 × 20 cm), defined as 9 central and 16 peripheral squares. At the beginning of the test, each animal was placed at the centre of the apparatus and then allowed to freely explore it for 5 mins. In between testing, the apparatus was cleaned using 70% ethyl alcohol to remove any odour cues from the previously tested animal.

We quantified parameters such as line crossing which is the total number of squares visited, rearing (the number of times the rat stood on its hind legs), grooming (the number of times the animal spent licking or scratching itself while stationary), freezing which is the duration with which the animal was completely stationary, centre square duration (the time the animal spent at the centre of the test box), stretched-attend Posture (the frequency with which the animal demonstrated forward elongation of the head and the shoulders followed by retraction to the original position) and defecation (the number of faecal boli).

##### The Elevated Plus Maze Test

The test is made of two wooden open arms (50 × 10 cm) crossed at right angles with two opposed arms of equal sizes as described by Walf and Frye [35]. Each animal is placed at the centre of the plus maze, facing one of the closed arms, and allowed to freely explore the arena for 5 min. The anxiety-related measures taken include the frequency and duration with which the animals visited the open arms and the closed arms.

##### Forced Swim Test (FST)

The forced swim test measures depressive-like behaviour in rodents [36]. For this experiment, each rat was placed in a cylindrical transparent container filled with enough water to keep its paws from reaching the base of the cylinder. As presumed by the test developer, the rats were supposed to make efforts to escape and after a while would adopt a posture of behaviour despair as indicated by immobility. The onset of immobility and the duration of immobility were recorded. The exposure duration was 6 minutes for each rat.

#### c. Tests for Spatial Learning and Memory

##### Morris Water Maze Test

The Morris water maze (MWM) behavioural test was employed for measuring the rats’ spatial learning [37]. A circular water pool with a hidden spherical escape platform was used in the experiment. The concealed platform is roughly two inches below the water’s surface. Each rat was to navigate its way to the platform using visual signals and the distal cue of a ribbon tied above the maze. The circular pool was labelled North, South, East, and West with the escape platform placed at roughly an equidistant angle between two cardinal points. Each rat was released into a different quadrant in the pool and subsequently to other quadrants to find the escape platform. The search duration was set at 120 seconds. Whenever the search for the platform was futile after 120 seconds, the rat would be guided to the platform and allowed to examine it for 15 seconds, then afterward, taken out of the circular pool entirely. This procedure was carried out on all the rats for five (5) successive days - marking it as the acquisition phase. The five (5) days acquisition phase of the Morris Water Maze was used to assess learning in the animal model.

The second phase of the assessment took place on the sixth day, and it involved the test of spatial memory. In this phase, the escape platform was removed from the pool, and then the rats’ memory was evaluated based on the time they spent within the quadrant where the platform was earlier placed. This evaluation also involved a converse cross-comparison of the time spent at the quadrant opposite the target quadrant. Evaluation and documentation of parameters of interest were done manually by two observers.

### Histopathology

#### Luxol Fast Blue (LFB) Staining

Brain sections (5 μm thick) were prepared and stained with Luxol Fast Blue (Electron Microscopy Sciences, Hatfield, PA, USA). Slides were kept in LFB solution and incubated at 58°C for 16 hours.

#### Measurements of inflammatory cytokines

Serum concentrations of IL-4, IL-10 and TNF-α were determined by commercially available high-sensitivity indirect sandwich enzyme-linked immunosorbent assay (Elabscience®). The preparation of all reagents, the working standards, and the protocol were followed according to the manufacturer’s instructions. The absorbance was read using an ELISA reader (BIO-RAD) at 450 nm and 570 nm dual filters. The detection ranges for TNF-α, and IL-10 were 7.8 pg/ml– 500 pg/ml, & IL-6 was 4.7 pg/ml- 300 pg/ml. All the sera samples were thawed once and assayed in duplicates.

### Data analysis

All behavioural test data were analyzed using a one-way analysis of variance (ANOVA) followed by Tukey’s post hoc test for comparison between groups. All tests were used with significance set at p < 0.05. The data were presented as mean ± standard error of the mean. One-way Analysis of Variance (ANOVA) was used to compare the mean levels of serum concentration between the control and Tukey post hoc test was used for multiple comparisons across group means. *P*-value < 0.05 was considered statistically significant.

## RESULTS

### Bisphenol-A does not alter forelimb strength, gross exploratory, and locomotor activities in Wistar rats

Although the control animals had the highest hanging latency in the forelimb support test, this latency was however not significantly different from the test groups (Figure 1). Additionally, we used the open-field test to evaluate locomotor activity in the experimental rats. In the present study the rearing, grooming, and stretched-attend posture frequencies, freezing, and number of fecal boli were similar across all experimental groups. Nonetheless, the BPA alone and BPA+ ascorbic acid groups had a statistically significant decrease in line crossing; (p= 0.0323), and (p=0.0389) respectively, when compared to the gallic acid alone group (Figure 2).

**Figure 1:**
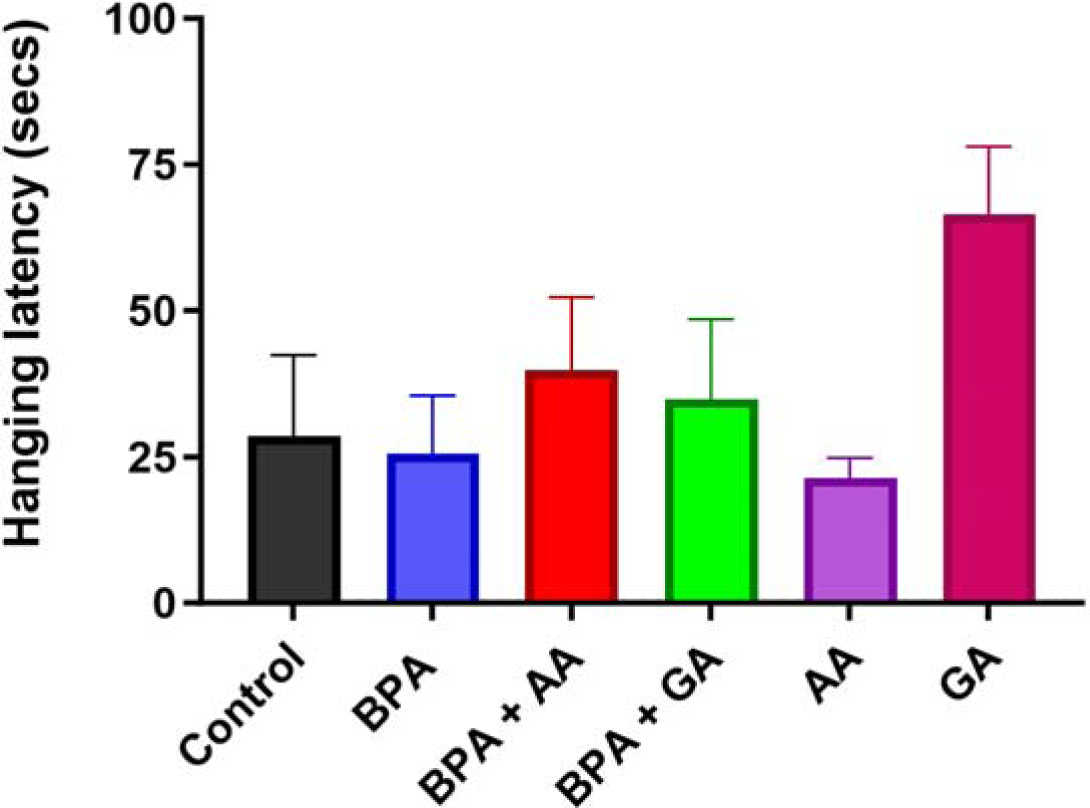
The effects of gallic acid (GA), ascorbic acid (AA), and BPA on forelimb grip test. Values are presented as mean + SEM (n=8/group). There was no significant difference in the hanging latency across the groups.

**Figure 2:**
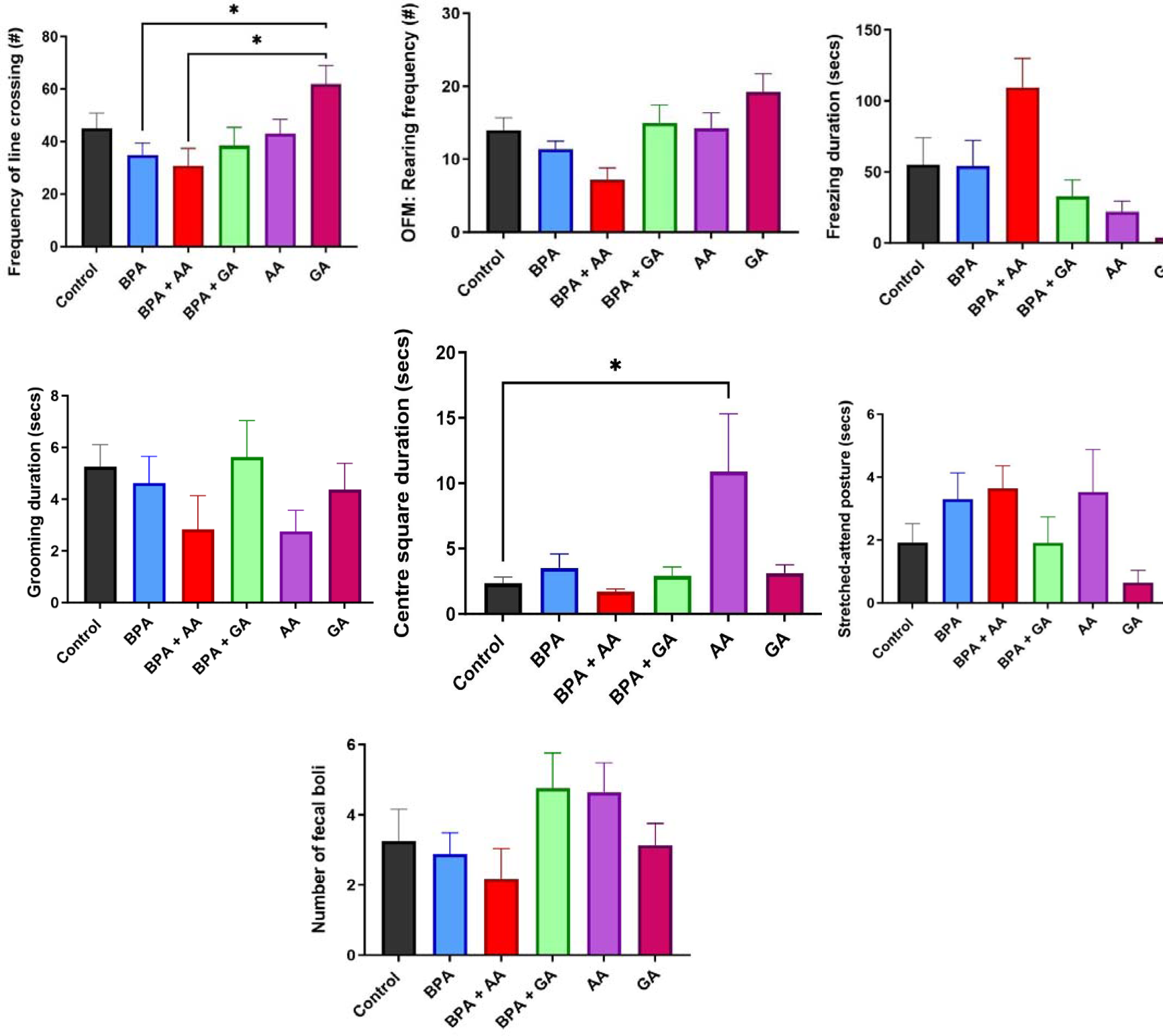
The effects of Gallic acid and Ascorbic acid on BPA-induced anxiety and locomotion, using the open field test (OFM). **A** is the number of line crossings; **B** is the rearing frequency; **C** is the grooming duration; **D** is the freezing time; **E** is the centre square duration; **F** is duration of stretched attend posture (SAP); **G** is the number of fecal boli. Values are presented as mean + SEM. There was no significant difference across all the groups in rearing, grooming duration, stretched attend posture (SAP), freezing, and fecal boli.

### Gallic and ascorbic acid ameliorated BPA-induced depressive-like behaviour in Wistar rats

There was a significant decrease in the time spent in the open arms in the BPA-alone group when compared to the GA-alone group (Figure 3A). Although the BPA alone group spent the least amount of time in the open arms, this was however not significantly significant when compared to the other experimental groups. Additionally, the duration in the closed arm of the elevated plus maze was significantly increased in the BPA alone, BPA+AA, and BPA+GA groups when compared with the other test groups. Furthermore, the BPA alone group spent more than 90% of the test duration (281.9 ± 31.76) in the closed arms. This prolonged duration in the closed arm in the BPA-alone group was significantly reduced in the control, AA-alone, and GA-alone groups (Figure 3D).

**Figure 3:**
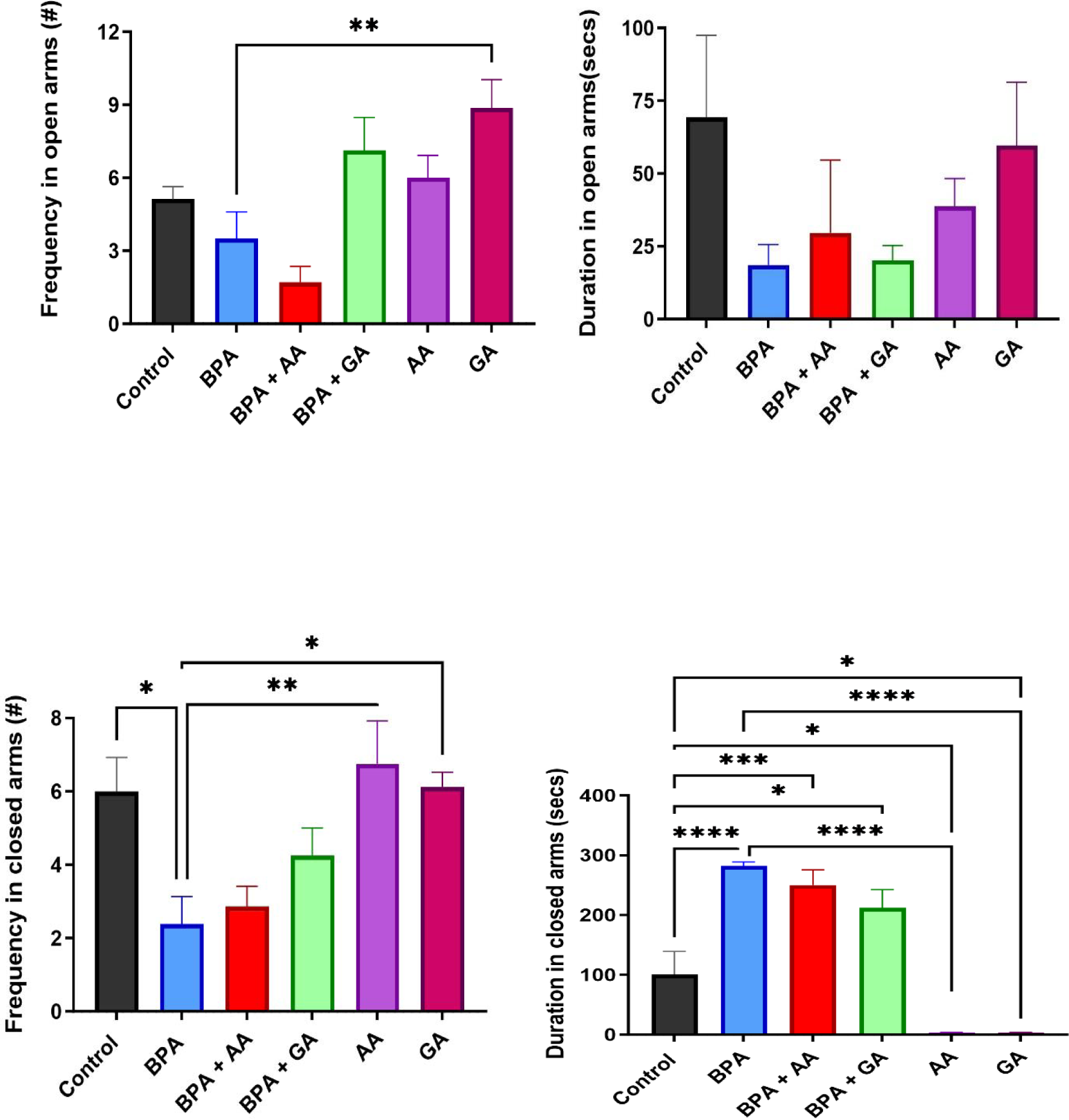
The effects of Gallic acid and Ascorbic acid on BPA-induced anxiety, using the elevated plus maze test. (**A)** number of entries into the open arms (**B)** time spent in the open arms (**C**) number of entries into the closed arms (**D)** time spent in the closed arms. Values are presented as mean + SEM. Superscript (a) indicates a significant difference at p < 0.05 compared with the control while superscript (b) Indicates a significant difference at p < 0.05 compared with the BPA group.

In the Forced swim test (FST), we observed the BPA group had the highest duration of immobility (143.2 ± 15.14) however, concurrent administration of BPA with gallic acid (BPA+GA group) led to a significant decrease in the duration of immobility ((Figure 4A). There was no significant difference in the onset of immobility across all the groups.

**Figure 4:**
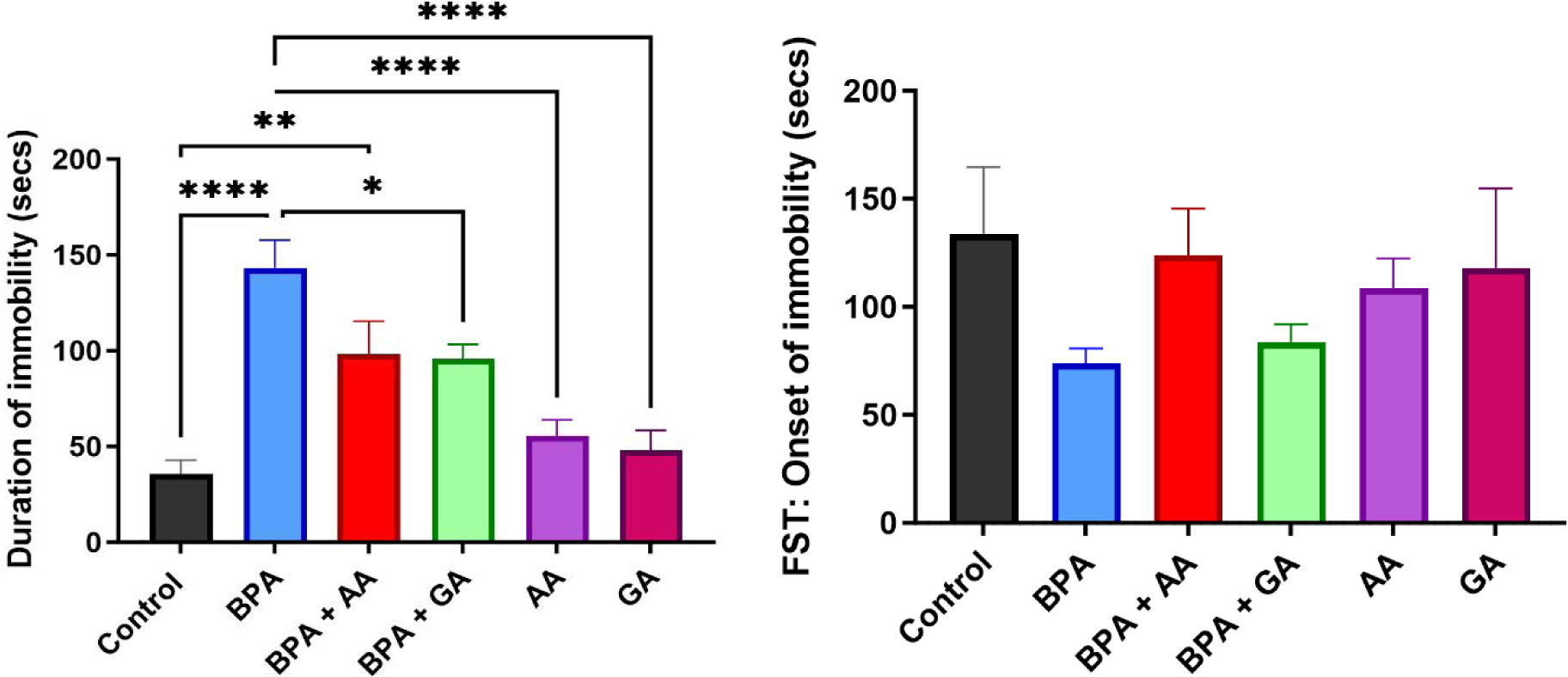
The effects of Gallic acid and ascorbic acid on BPA-induced depression in the forced swim test. **A** is the duration of immobility. **B** is the onset of immobility. Values are presented as mean + SEM. Superscript (a) indicates a significant difference at p < 0.05 compared with control (Group A), while superscript (b) Indicates a significant difference at p < 0.05 compared with the BPA alone group.

**Figure 5:**
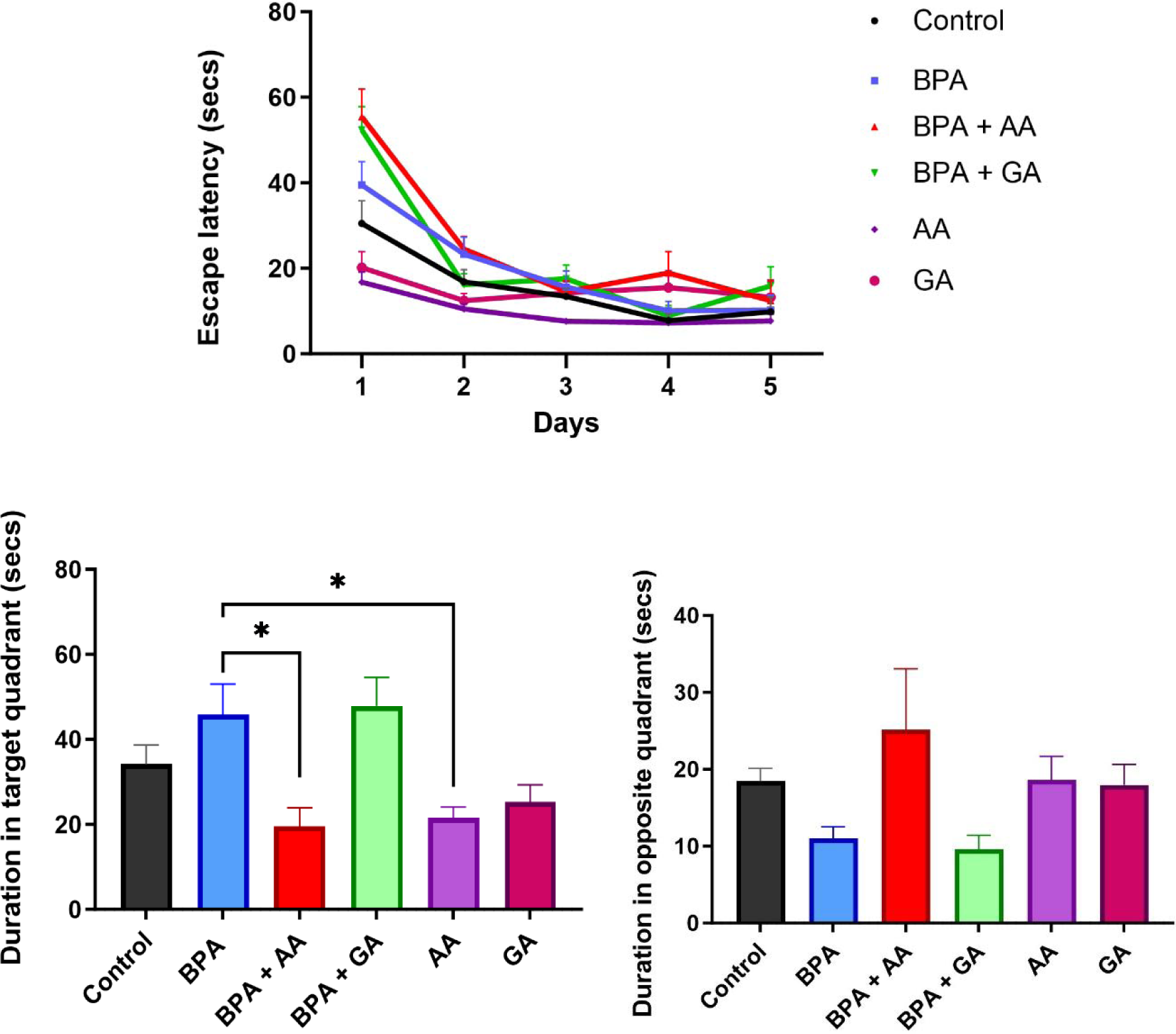
**A.** Acquisition phase of Morris water maze test for 5 days. **B**. The time spent in the target quadrant. **C**. The time spent in the opposite quadrant. Values are presented as mean ± SEM. Superscript (a) indicates a significant difference at p < 0.05 compared with control (Group A), while superscript (b) Indicates a significant difference at p < 0.05 compared with Group B.

**Figure 6:**
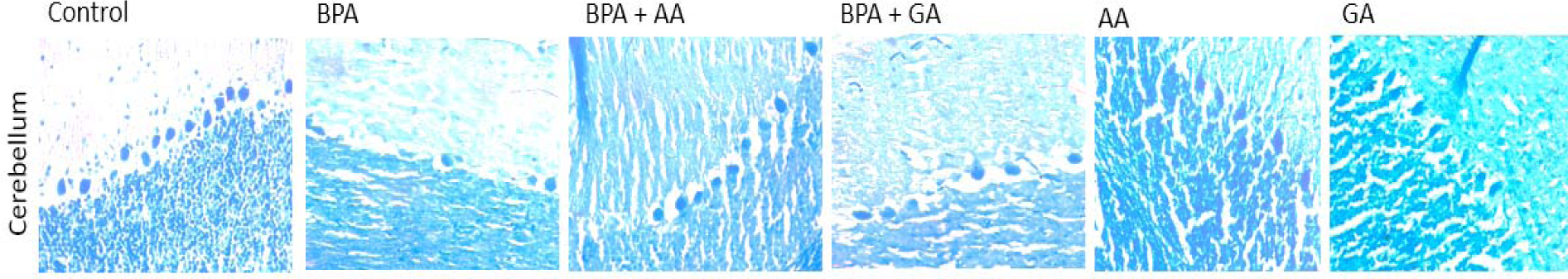
Luxol fast blue staining of the cerebellum. The figure shows the Purkinje cell layer of the cerebellum. The BPA group showed a sparse Purkinje cell layer compared with the control group. There was restoration of this layer in both groups treated with GA and AA.

**Figure 7:**
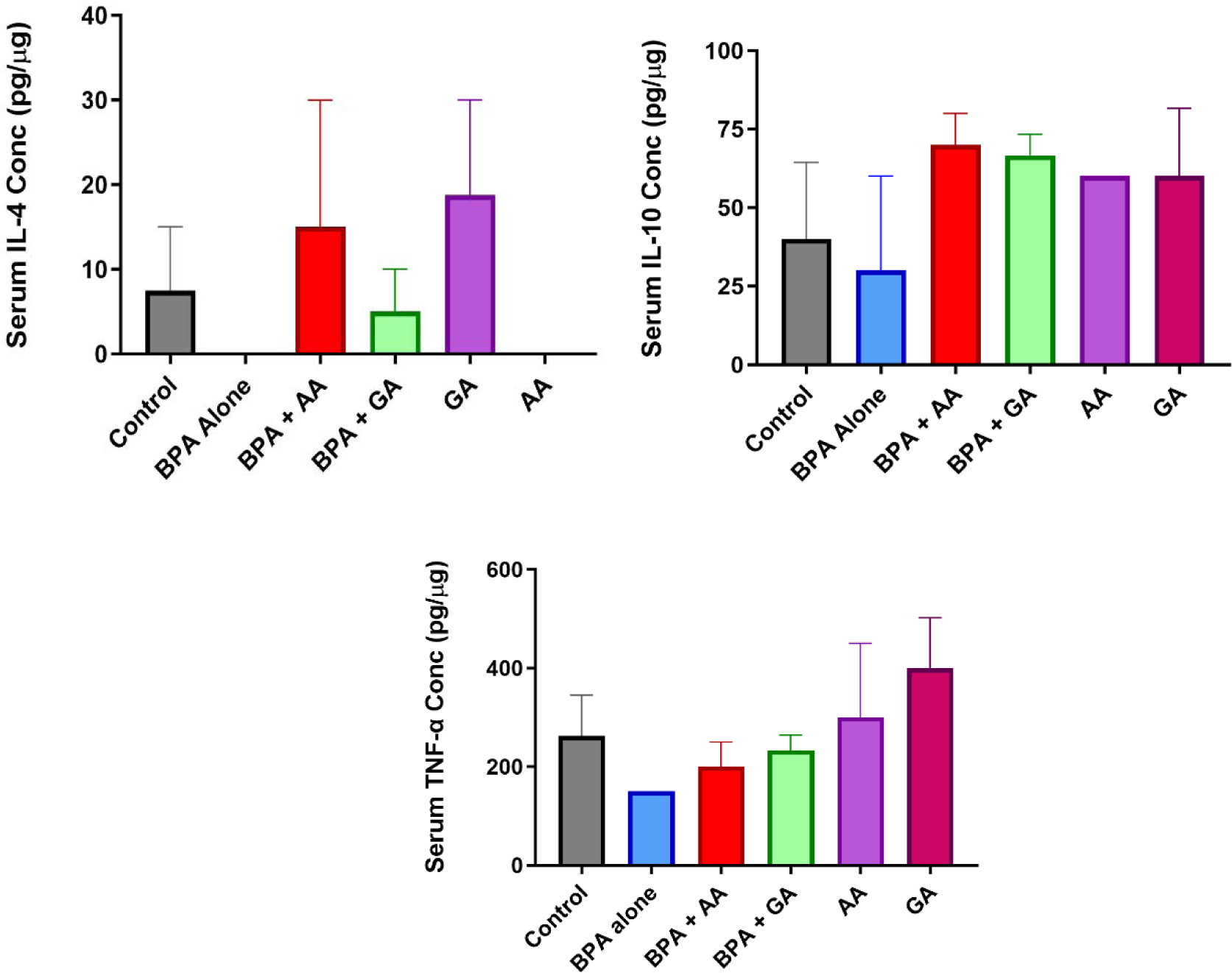
The levels of inflammatory cytokines IL-4, IL-10, and TNF-alpha in the serum. There was no significant difference in the levels of inflammatory cytokines across all the groups.

### Learning and spatial memory were intact in experimental rats

During the acquisition phase of the Morris water maze test, we observed a decrease in the escape latency following an increasing number of trials across all the groups. This implies the animals learn the rules of the test across all the groups and spend reduced time exploring the hidden platform in the maze as they do more trials in the first phase of the test. Similarly, during the probe phase of the test which is a test for spatial memory the duration in the target quadrant was similar in the BPA alone group compared to the control. However, the BPA+AA and AA alone groups spent significantly less time in the target quadrant compared to the BPA alone groups.

### Brain histology staining for myelin revealed cerebellar demyelination in the BPA rats

Myelinated regions in the cerebellum were visualized with the Luxol Fast Blue staining technique. Highly myelinated areas were stained dense whereas demyelinated or hypomyelinated areas were stained pale. Demyelination was not observed in brain sections of control, AA, and GA groups. However, the cerebellar sections of BPA rats appeared pale. Similarly, we observed a loss of Purkinje cells in the BPA-alone groups.

### Levels of inflammatory cytokines

Due to previously documented increased generation of reactive oxygen species (ROS) following BPA administration, we evaluated the downstream effects on cytokines production. We observed a decrease in serum levels of TNF-α and IL-10 in the BPA group. In addition, IL-4 was not detected in the serum of mice in the BPA-alone and AA-alone groups.

## DISCUSSION

BPA has been reported by several studies to induce aggression, anxiety, cognition, and learning-memory deficits [38]. One of its mechanisms is the generation of reactive oxygen species (ROS) and reactive nitrogen species (RNS) which leads to oxidative stress [39]. Gallic acid is a documented antioxidant with neuroprotective properties [30]. Ola-Davies and Olukole [29] reported gallic acid protects against BPA-induced cardio-renal alterations through the antioxidant defense system.

In the present study, we evaluated the behavioural alterations associated with a sub-chronic (21 days) administration of bisphenol-A. Besides, we assessed the effects of simultaneous administration of gallic acid (GA) and ascorbic acid (AA) after giving BPA in mitigating some of these behavioral impairments.

First, we examined the forelimb motor strength using the hanging wire test, and our findings revealed that the hanging latency was similar across all groups indicating that Bisphenol-A did not alter forelimb grip strength. Our results are in congruence with previous data by Miyagawa et al., [16] and Ji et al., [41], who did not detect any motor impairment in mice postnatally exposed to BPA. Findings in the literature are rare on the effects of bisphenol-A toxicity on grip strength, hence there is a need for further studies to validate or refute our present findings.

Regarding locomotion, we observed that BPA did not change the total exploratory time compared to control rats, although administration of BPA induced a reduction in exploration compared to the GA alone group. Furthermore, we used the elevated-plus maze to analyze anxiety-related behaviour based on the hypothesis that there is extreme stress from being in the open arms versus the closed arms of an elevated maze. We observed anxiogenic responses in the BPA group as revealed by a reduction in duration and number of entries into the open arm. This anxiety-like activity of BPA has been observed previously [42, 43]. The observed anxiogenic effect following the administration has been linked to the downregulation of the alpha-1D adrenergic receptor in the paraventricular thalamus by BPA [41] and changes in gene expression associated with affective behaviours [44]. However, the concomitant administration of gallic acid mitigated the anxiety-like behaviour of BPA.

BPA has also been reported to cause depressive-like behavior [40]. Depression is evaluated in rodents with the forced swim test as a model [45]. The duration of immobility is a measure of depression [36]. The increased duration of immobility seen in the BPA-alone group corroborates the report by van den Bosch and Meyer-Linderberg, (2019) [40]. Only GA ameliorated this effect as seen by the reduced duration of immobility seen in the BPA and GA-treated group as there was no significant difference in AA treated group.

The Morris water maze is a test for learning and spatial memory [37]. It tests for hippocampal-dependent learning, which includes acquisition and long-term memory [46]. In the present study, learning occurred across all the groups as the time taken to reach the target quadrant reduced although there was only a significant difference between the BPA group treated with GA and the control group. There was no difference in the probe trial as a test for spatial memory in the experimental rats. These findings are consistent with the reports documented by Bowman [47] who reported that learning and spatial memory were intact following exposure of adolescent rats to BPA. Still, some authors reported impairment in learning and spatial memory following the administration of BPA in experimental rodents [48, 49]. The incongruence in our findings with these reports may be explained by the difference in the duration and dosage of administration of BPA. We hypothesize that this lack of impairment in learning and spatial memory in our study may be due to the shorter periods of administration of the toxicant in the present study.

Since oxidative stress has been associated with triggering the secretion of some pro-inflammatory cytokines, we thus evaluated the levels of IL-4, IL-10, and TNF-α in the serum. Interleukin-4 and Interleukin-10 are pleiotropic cytokines that act predominantly to suppress the pro-inflammatory environment by downregulating the expression of inflammatory cytokines such as TNF-α, IL-6, and IL-1 by activated macrophages. In addition, IL-10 can up-regulate endogenous anti-cytokines and down-regulate pro-inflammatory cytokine receptors. Thus, it can counter-regulate the production and function of pro-inflammatory cytokines at different levels. However, there was a reduction in the serum level of these anti-inflammatory cytokines following BPA administration which is indicative of its inflammatory potentials. It has been previously reported that administration of BPA stimulates the synthesis of pro-inflammatory cytokines, such as and interleukin (IL)-6 and tumor necrosis factor-α (TNF-α) but inhibits synthesis of anti-inflammatory cytokines, such as IL-10 in cellular studies via ER/nuclear factor- κB (NF-κB) signaling pathway [50] (Liu et al., 2014). In the present study, concomitant administration of gallic acid albeit led to the decreased production of pro-inflammatory cytokines to level similar with control.

The behavioral disruptions of BPA reported in our study may be due to its effects on brain regions, such as the cerebellum, and several other areas such as the nucleus accumbens, thalamus, and amygdala as earlier reported [51, 52, 53]. We are thus currently investigating further studies to examine different brain regions outside of the cerebellum. This will help us to further elucidate the mechanisms involved in the alteration of behaviour by BPA.

## CONCLUSION

Our results show that the primary effect of bisphenol-A toxicity-induced behavioural impairment behaviors is anxiety and depressive-like rather than memory and locomotor activity impairments. Gallic acid ameliorates behavioural deficits induced by BPA, thereby it can be potentially used in the treatment of neurological disorders associated with BPA and other environmental toxicants. Further investigation will however be needed to understand the neurobiological mechanisms underlying the ameliorative effects of GA on BPA with biochemical analysis, histology, and immunohistochemistry.

## List of Abbreviations

BPA: Bisphenol-A
AA: Ascorbic acid
GA: Gallic acid
LFB: Luxol fast blue
FST: Forced Swim Test
MWM: Morris Water Maze

